# BiomiX, a User-Friendly Bioinformatic Tool for Automatized Multiomics Data Analysis and Integration

**DOI:** 10.1101/2024.06.14.599059

**Authors:** Cristian Iperi, Álvaro Fernández-Ochoa, Guillermo Barturen, Jacques-Olivier Pers, Nathan Foulquier, Eleonore Bettacchioli, Marta Alarcón-Riquelme, PRECISESADS Flow Cytometry Study Group, PRECISESADS Clinical Consortium, Divi Cornec, Anne Bordron, Christophe Jamin

## Abstract

BiomiX addresses the data analysis bottleneck in high-throughput omics technologies, enabling the efficient, integrated analysis of multiomics data obtained from two cohorts. BiomiX incorporates diverse omics data. DESeq2/Limma packages analyze transcriptomics data, while statistical tests determine metabolomics peaks. The metabolomics annotation uses the mass-to-charge ratio in the CEU Mass Mediator database and fragmentation spectra in the TidyMass package while Methylomics analysis is performed using the ChAMP R package. Multiomics Factor Analysis (MOFA) integration and interpretation identifies common sources of variations among omics. BiomiX provides comprehensive outputs, including statistics and report figures, also integrating EnrichR and GSEA for biological process exploration. Subgroup analysis based on user gene panels enhances comparisons. BiomiX implements MOFA automatically, selecting the optimal MOFA model to discriminate the two cohorts being compared while providing interpretation tools for the discriminant MOFA factors. The interpretation relies on innovative bibliography research on Pubmed, which provides the articles most related to the discriminant factor contributors. The interpretation is also supported by clinical data correlation with the discriminant MOFA factors and pathways analyses of the top factor contributors. The integration of single and multi-omics analysis in a standalone tool, together with the implementation of MOFA and its interpretability by literature, constitute a step forward in the multi-omics landscape in line with the FAIR data principles. The wide parameter choice grants a personalized analysis at each level based on the user requirements. BiomiX is a user-friendly R-based tool compatible with various operating systems that aims to democratize multiomics analysis for bioinformatics non-experts.

**Key points:**

- BiomiX is the first user-friendly multiomics tool to perform single omics analysis for transcriptomics, metabolomics and methylomics and their data integration by MOFA in the same platform.
- MOFA algorithm was made accessible to non-bioinformaticians and improved to select the best model automatically, testing the MOFA factor’s performance in groups separation.
- Large improvement of MOFA factor’s interpretability by correlation, pathways analysis and innovative bibliography research.
- BiomiX is embedded in a network of other online tools as GSEA, metaboanalyst EnrichR etc, to provide a format compatible with further analyses in these tools.
- Interface and usage are intuitive and compatible with all the main operating systems, and rich parameters are set to grant personalized analysis based on the user’s needs.

---

The explosion of high-throughput technologies has enabled the generation of vast amounts of data on multiple levels of biological organization, as observed in the autoimmunity field with the European PRECISESADS database [1, 2], which includes groups of patients with seven autoimmune diseases and controls. This revolution brought new tools for analyzing single omics with high efficiency. The most common packages are Deseq2 [3], EdgeR [4], and Limma [5]. for transcriptomics RNA sequencing while ChAMP [6] and IMA [7] R are packages for methylomics analysis. For metabolomics, MetID [8] is an R tool that connects many functionalities to work but requires bioinformatics knowledge while Metaboanalyst’s [9] was noteworthy for users without a bioinformatics-metabolomics background but has a limited annotation customization. During the last decade, a significant boost in omics integration arised. The state of the art includes many strategies based on diversified algorithms and models, including matrix factorization– regression and association methods such as MOFA [10] and DIABLO [11], other matrix factorization methods [12,13], IclusterPlus [14], and network analysis. These include Bayesian networks such as PARADIGM [15] and matrix factorization-based methods such as NEMO [16] and SNF [17]. However, each available tool was developed to solve a specific task, such as disease subtyping, disease insight, or biomarker prediction. These tools demand expertise in coding and bioinformatics, and unfortunately they often remain far from the critical eye of specialized biologists and clinicians. The interest in multiomics data is supported by databases containing cross-analysis on multi-omics, as Cancer Genome Atlas (https://cancergenome.nih.gov/), and the Omics Discovery Index (https://www.omicsdi.org). The datasets available in single-omics repositories include Gene Expression Omnibus (https://www.ncbi.nlm.nih.gov/geo/) for transcriptomics and methylation, Proteomics Identification Database PRIDE (https://www.ebi.ac.uk/pride/) for proteomics [18], and MetaboLights (https://www.ebi.ac.uk/metabolights/) for metabolomics data [19]. Bioinformaticians and data scientists must grant access to novel bioinformatics sources and tools, notably specialists in their fields. This need has driven the development of BiomiX, a solution to ease access for users without bioinformatics expertise and to allow them to focus on data interpretation. BiomiX provides robust, validated pipelines in single omics with additional functions, such as sample subgrouping analysis, gene ontology, annotation, and summary figures. Moreover, to our knowledge, BiomiX is the first bioinformatics tool to simultaneously include single-omics analyses and their integration. BiomiX implements MOFA, allowing for an automatic selection of the total number of factors and the identification of the biological processes behind the factors of interest through clinical data correlation and pathway analysis. BiomiX implemented, for the first time, the factor identification through an automatic bibliography research on Pubmed, underlining the importance of integrating literature knowledge in the interpretation of MOFA factors. The graphic user interface of BiomiX is available to ensure user-friendliness and flexibility and handle transcriptomics, metabolomics, and methylomics data.

## Program Description and Methods

### Graphical user interface and R environment

BiomiX interface was developed using the Python toolkit PyQt5. It allows to choose the analysis and desired parameters for each omics (transcriptomics, metabolomics, and methylomics) to provide the output results and the normalized data required for integration analysis. The global script of BiomiX system is shown in **Figure 1**. The interface is available on all OS systems, such as Windows, Linux, and Mac. The download and tutorial are available on the following BiomiX github pages, respectively: https://github.com/IxI-97/BiomiX2.2 and https://ixi-97.github.io. The installation occurs by conda environment.

**Figure 1.**
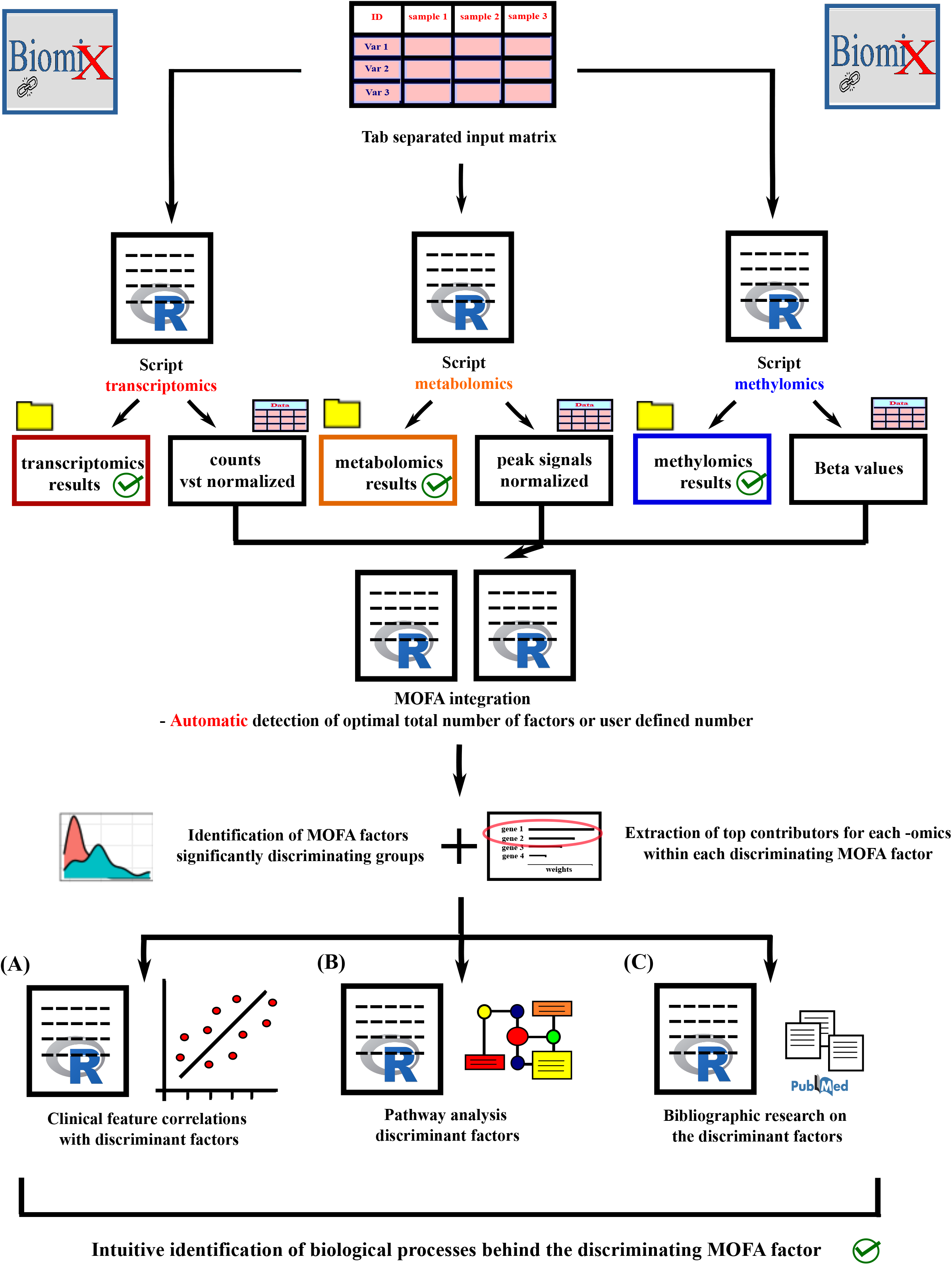
BiomiX Pipeline and Script Schema. Illustration of the scripts and the general BiomiX structure. The upper part shows the input table and the three main scripts for analyzing single omics, including the output results and normalized matrixes. These matrixes are used as inputs for MOFA integration. The user can automatically optimize or arbitrarily select the total number of factors. In both cases, a distinct script is running. The bottom part represents the definition of the discriminant factor and the extraction of its top features. The discriminant factors are correlated with clinical data to spot significant correlations using Pearson correlation (A), while the top features are explored by pathway analysis (B) and a text-mining/PubMed research approach (C).

*BiomiX interface parameters* – BiomiX aims to provide a simple, intuitive user interface, as shown in **Figure 2**. It first asks users for a metadata file containing samples to analyze in the omics databases. BiomiX automatically generates the main interface, which allows users to select a detected group as a control (CTRL) and a condition/disease group for analysis. It displays any group in the provided database. The interface has six rows, representing slots for omics data. Each column allows users to better define the input and the analysis to be performed. It includes a prompt to specify whether the data must be analyzed or integrated and asks for the type of omics and the label to name the output folder. This label can also generate a regex to filter samples using sample names. Finally, an input button allows the uploading of the matrix file to BiomiX. Here users are asked if they wish to modify the matrix format. If the answer is affirmative, an assisted format converter guides users through the conversion process to the BiomiX format. Any transcriptomics, metabolomics and methylomics data can be added, analyzed, and integrated into any combination desired by users. Data integration is available without repeating the single-omics analysis when the first single-omics analysis is completed, as the normalized data are saved in the MOFA input and ready to be integrated. Once the databases are selected for integration, the parameters for MOFA integration in the lower section of the interface and advanced options are established. Users are able to define an arbitrary number of MOFA factors or set an automatic selection of the best total number of factors in the MOFA model. Users also could select one factor from the interface to focus on in the final report, in which its omics contributions, clustering and the heatmap are displayed. MOFA allows the integration of samples that are not shared in all the omics, but it could lead to error if this is the case for most of the database. Therefore, BiomiX adds a parameter to filter the samples in the integration analysis using a minimum number of shared omics.

**Figure 2.**
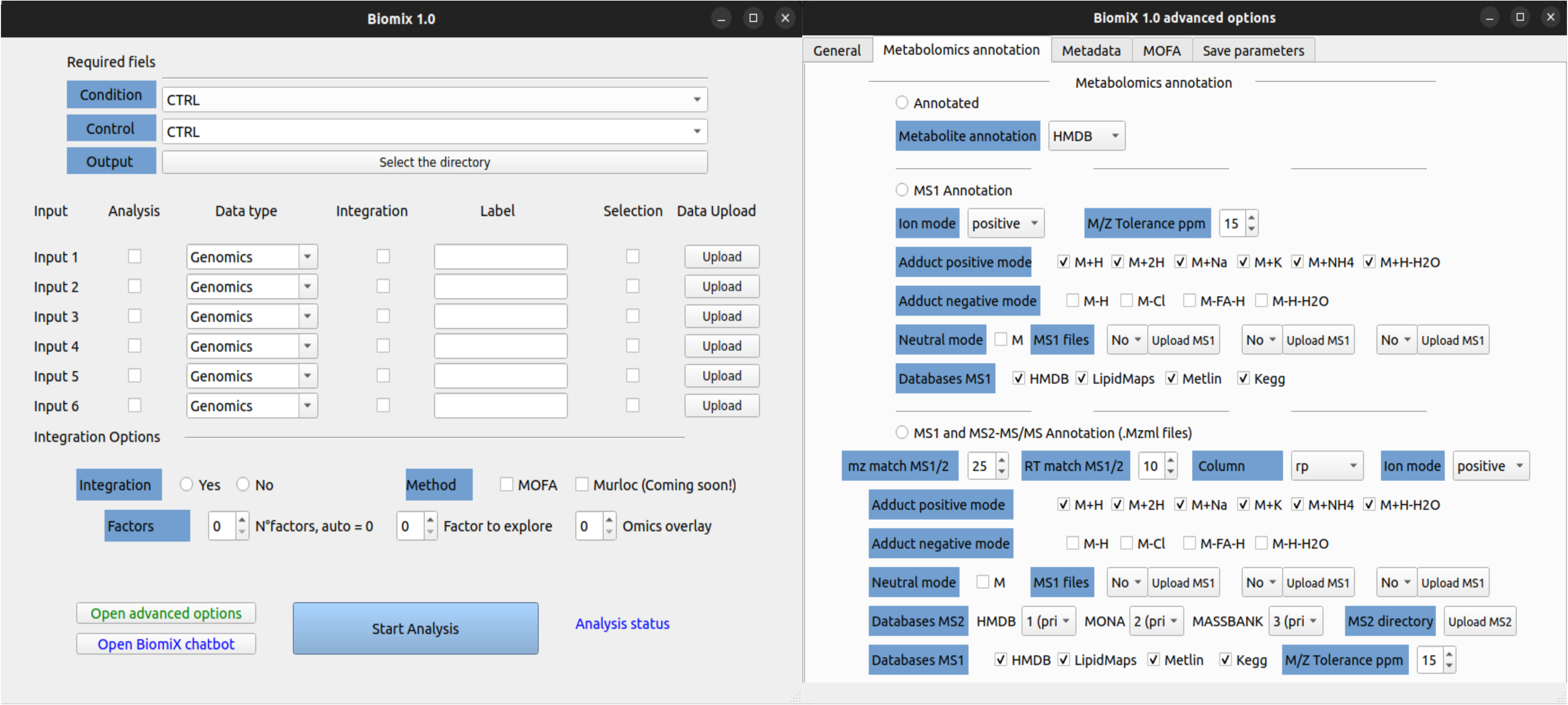
BiomiX Interface. Illustration of the BiomiX main interface on the left and advanced options windows on the right. The advanced options are divided into four sections: general, metabolomics annotation, metadata, and MOFA.

Advanced options allow for deeper analysis customization and can be divided into five sections. The first is the “general” section, which includes Log2FC and the adjusted p-value threshold, CPU usage, number of input variables for MOFA, gene panel, and Panel positivity criteria. The second section is the metabolomics section, which allows users to select the desired annotation type, including none, MS1 and MS2. The MS1 files, containing mass-to-charge ratio (m/z) and retention time, could be uploaded in this section as the directory for the mzML files for MS2 annotation. The prioritization of the available metabolomics databases, such as HMDB, KEGG, LipidMap, Metlin, MassBank, and Mass Bank of North America (MoNA) could then be established. The third section allows users to filter the samples based on the metadata information provided, where it is possible to filter on a threshold or a group in a selected metadata column (e.g. cell purity, ethnicity and proteinemia). The fourth section allows for customizable MOFA analysis, setting speed and iteration of the model and threshold in contribution weights. It also affects MOFA interpretation, including the number of articles considered in the bibliography research, the type of clinical data available, and the p-value threshold in pathway analysis. The last section allows you to save the parameters selected.

### The BiomiX-assisted format convertor

This is a simple functionality that allows users to modify a matrix directly in the BiomiX interface. It can also perform transposition, remove columns or rows and identify the features column. It is designed to facilitate the conversion of any data table to the BiomiX format.

### BiomiX transcriptomics input and pipeline

BiomiX requires the expression matrix M_sg_ where the columns “**s**” represent the samples, and the rows g contain the genes in Ensembl or the gene name symbol. The matrix must contain the row counts in integer format obtained by counting the aligned reads in the genes in the bam files. Examples of tools are HTSeq-count [20] or featureCounts [21]. Alternatively, normalized data can be used as input in floating format. Limma automatically recognizes and analyzes them. If sex and gender are available in the metadata, they are automatically used to correct the Deseq2 and Limma models. The row counts are used as input for the Deseq2 and Limma R packages to compare the two conditions defined in the “condition” column of the metadata to conduct differential gene expression analysis. Differentially expressed genes are then sorted in the results files and separated by their significance and up- or down-regulation compared to the CTRLs. The default thresholds to consider the differentially expressed genes are set to Log2FC > |0.5| and p.adj < 0.05. The p-value is adjusted using the FDR method. A heatmap and a volcano plot with increased and decreased top genes are automatically created. Users can choose the number of visualized genes. The enrichment of biological processes in the results is explored in the R version of EnrichR. Moreover, output files are produced as input for gene set enrichment analysis using Gene set enrichment analysis (GSEA) [22] (http://www.gsea-msigdb.org/gsea/index.jsp) or the EnrichR web tool [23] (https://maayanlab.cloud/Enrichr/). The expression matrix is also normalized by variance-stabilizing transformation for data visualization and MOFA integration. If a gene panel is uploaded, the same analysis and output files are generated for the negative subgroup versus the CTRL group and the positive subgroup versus the CTRL group. This can confirm subgroups or define new subpopulations within known ones for diseases or treatments with well-known subpopulation markers (*e.g.* interferon or interleukin signaling genes in autoimmune diseases…). A summary of these features and the pipeline is shown in **Figure 3**.

**Figure 3.**
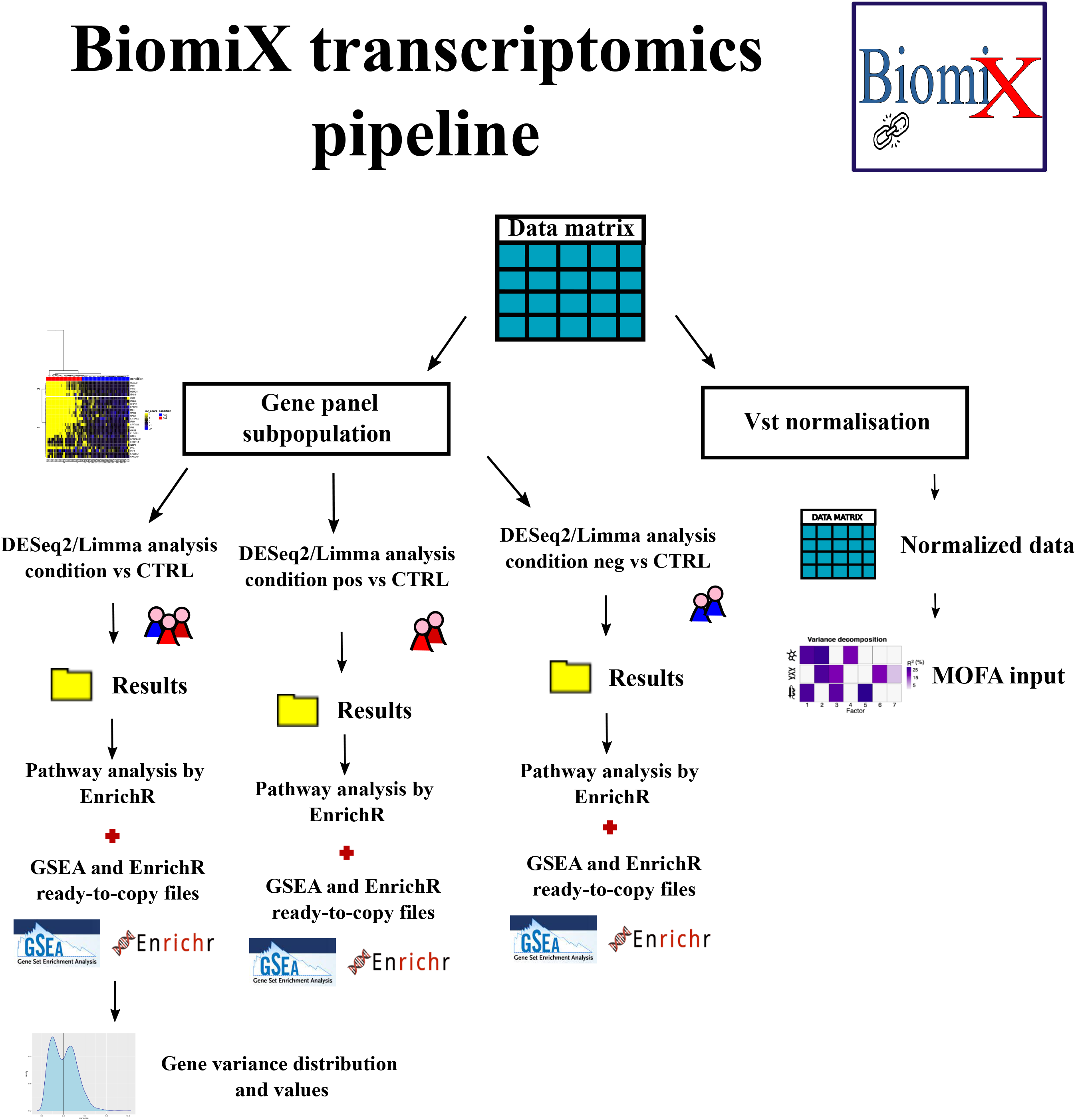
BiomiX Transcriptomics Pipeline. Illustration of the BiomiX transcriptomics pipeline, showing the generation of normalized data via Vst normalization for MOFA integration on the right and statistical analysis on the left. Statistical analysis includes a subpopulation step in which additional differential gene expression analysis by DESeq2 or Limma is performed on subpopulations using the control (CTRL) as a reference. A positive (pos) and negative (neg) subgroup are created. For the subpopulation, a gene panel file must be provided. The results folders generated in the analysis (pos vs CTRL and neg vs CTRL) include the volcano plot and the results table, including Log2FC, p-value, and False Discovery Rate (FDR). The up-and down-regulated genes are also separated to ease their exploration and to copy them to EnrichR for pathway analysis. A preliminary analysis using EnrichR on R is already available in the results. The expression matrix is also converted into .gct format as input for GSEA analysis. The gene variance distribution is explored, providing plots and value.

### Subpopulation of differential gene expression analysis (DGE) based on a gene panel

For transcriptomic analysis, BiomiX enables the insertion of a panel of genes to identify subpopulations within the condition being analyzed to conduct separate DGE analyses. It compares positive and negative patients for the gene panel with the CTRL group, providing the same results files for any transcriptomic analyses. Subpopulation recognition relies on the variation measured in standard deviation units compared to the mean expression in the chosen CTRL. This method was inspired by similar approaches to measuring IFN-alpha signaling, such as the Kirou score [24] or similar methodologies [25]. Any panel of genes of interest could be similarly employed. Specifically, the standard deviation score (Z activity score) uses the counts normalized by the number of reads to compare the expression of each gene (g) in each disease or treated sample (s) with the mean expression of the CTRLs divided by the standard deviation of the CTRLs as in the following equation:

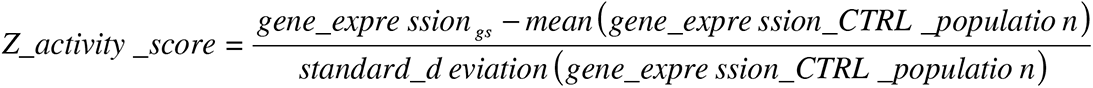

The higher the standard deviation shift is in a gene, the higher the gene expression is in the condition compared to the CTRLs. Subgrouping is dependent on the user parameters and criteria and is completely customizable. By default, according to the Kirou score, the samples with three genes with a score > 2 or 10 genes with a score > 1 are labeled positive. A heatmap is built using the standard deviation score, based on hierarchical clustering in the Complexheatmap package v2.12.022 on R. Euclidean distance, and Ward’s D2 method is used for hierarchical clustering by default.

### BiomiX metabolomics input and pipeline

BiomiX metabolomics data requires a peak signal matrix M_sg_, where the columns “**s**” represent the samples, and the rows “**g**” contain the arbitrary peak numbers. BiomiX is flexible enabling both targeted and non-targeted metabolomics data to be obtained. Untargeted data allow for annotated and non-annotated metabolomics peaks. It can provide annotation if required based on MS1 annotation (mass/charge ratio “m/z” and retention time values) or MS2 data (MS1 annotation plus raw MS2 fragmentation files in mzML format). Users starting with mzML files can generate the peak matrix by pre-processing raw data (peak deconvolution, RT alignment, and normalization) by user-friendly tools (MZMine, MS-DIAL and Metaboanalyst) or R packages pipelines [26–28]. Examples of these tools can be found in Á. Fernández-Ochoa et al.’s or X. Shen’s articles [29,30].

The peak signals from the treated samples are compared with those of the CTRL samples, calculating the Log2FC as the log2 of the ratio between the median peak signal in the condition-treated samples and the median peak signal in the CTRL samples for each peak. The p-values are evaluated by the non-parametric Mann–Whitney test and corrected through the FDR method. Then, the metabolomics peaks are annotated using the CEU Mass Mediator tool [31] through the CMMR R package [32], which provides a direct application programming interface in R. The MS1 m/z match is set by default to a 15-ppm error for positive mode, but neutral and negative modes are also available. The adducts available in the positive mode include M+H, H+2H, H+NA, H+NH4, and M+H-H2O, and in the negative mode, they include M-H, M-Cl, M+FA-H, and M-H-H2O. By default, all available MS1 databases are examined (the Human Metabolome Database [HBMD], Lipidmaps, Metlin, and Kegg), but their use is customizable. BiomiX automatically examine the lists containing previously identified or predicted metabolites in the HBMD [33] to filter metabolites at the same time associated with one identical peak for retaining those already identified or spotted in a type of specimen. These include plasma, urine, saliva, cerebrospinal fluid, feces, sweat, breast milk, bile and amniotic fluid samples. When MS/MS spectra are also available, a metabolite annotation based on the fragmentation spectra is also possible. BiomiX will automatically upload all the mzML files in the indicated directory and verify the peak fragmentation spectra, looking for a match in the Mass Bank of North America (MoNA), MassBank, and HMDB. The user must prioritize these databases, using the first as a reference and the others to fill the metabolomics peaks not annotated by the higher-priority databases. Priority and use of these databases are fully customizable, but the default order of priority is HMDB, MoNA, and MassBank. The overlap of the candidate spectra retrieved in mzML files and those from the databases are saved in the output folder. Each peak annotation detected in MS2 will automatically replace the annotation obtained in MS1 because the former is more reliable. A summary of these features and the pipeline are shown in **Figure 4**.

**Figure 4.**
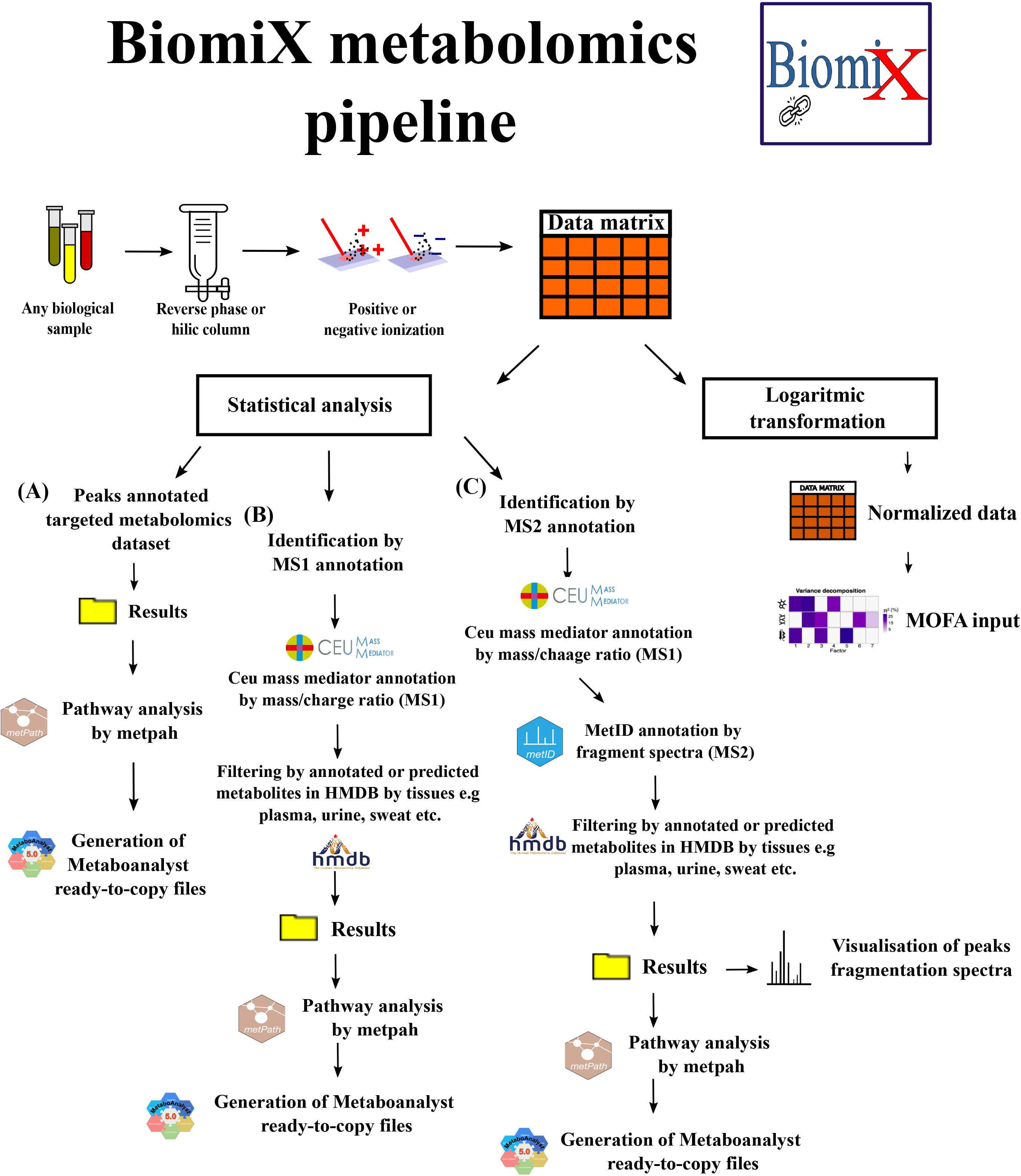
BiomiX Metabolomics Pipeline. Illustration of BiomiX metabolomic pipeline after ionization of samples, showing the generation of log-transformed data for MOFA integration on the right and statistical analysis on the left. The statistical analysis includes three pipelines based on the peak annotation status. If the annotation is already present (pipeline “A”), the metabolomic peaks among the two conditions (disease and CTRL) are directly analyzed to obtain the results folder including Log2FC, p-value, and false discovery rate (FDR). MetPath analyzes the biological pathways of significant metabolites and treats them as ready to copy on Metaboanalyst for further analysis. If transcriptomics data are available, BiomiX produces input files for Metaboanalyst joint pathway and network analyses. The metabolomics annotation based on the MS1 information (m/z) (pipeline “B”) is available. Here, the pipeline exploits the CEU Mass Mediator database to match the m/z of the metabolomics peaks with those provided by the database, reporting the best matches for each peak. As one m/z value can have multiple annotations, the candidate can be filtered based on reports or predictions of that metabolite in the sample type (i.e., human plasma, urine, feces, and saliva) by HMDB. The remaining pipeline includes statistical and pathway analyses using MetPath and Metaboanalyst, as described previously. The metabolomics annotation of the fragmentation spectra (MS2) together with MS1 information (pipeline “C”) is similar to pipeline “B,” adding an exploratory analysis on the fragmentation files (mzML) after the CEU Mass Mediator annotation. Here, HMDB, MassBank, and MoNA databases are examined to match each fragmentation spectra in the sample with those available in these databases. These matches are available in the results folder. The annotation based on the fragmentation spectra MS/MS (MS2) retrieved in this step will replace those obtained based on the MS1 annotation because of its higher reliability. As described previously, the remaining pipeline includes statistical and pathway analyses using MetPath and Metaboanalyst.

The top increased and reduced significant metabolites are displayed in a volcano plot and heatmap, while a logarithmic transformation is applied to the peak signal matrix for MOFA integration. The user chooses the number of metabolomic peaks displayed. To unveil the change in biological pathways, BiomiX automatically exploits the R packages MetPath v1.0.5 from TidyMass v1.0.8 [29] to spot enrichment in the metabolic process among the significantly changed metabolites. Moreover, a deeper analysis prepares ready-to-copy files as input for Metaboanalyst [9]. It is one of the most advanced online tools for metabolomics analysis, including enrichment and metabolite set enrichment analysis. When transcriptomics data are available in BiomiX analysis, metabolomic and transcriptomic results are autonomously integrated into copiable files on Metaboanalyst to launch joint pathway and network analyses.

### BiomiX methylomics input and pipeline

BiomiX requires the expression matrix “M_sg_,” where the columns “**s**” represent the samples, and the rows “**g**” contain CpG island annotation. The matrix must contain beta values; if unavailable, these can be obtained using the Minfi R package [34]. Differential methylation analysis is performed using the ChAMP [6] database, providing the results of the CpG island hypermethylated and hypomethylated with a volcano plot. The threshold has been set as the default to the beta value change (Δbeta) > |0.15| and p.adj < 0.05, but the user can customize it. Each methylomics analysis provided a volcan^Δ^o plot containing the names of the top CpG islands with increased and reduced methylation, as well as a heatmap including the top CpG islands with increased and reduced methylation between the two conditions. The users chose the number of CpG islands to visualize. A complete list of CpG islands with increased or reduced methylation is created. Each CpG is associated with the gene, chromosome, Log2FC, adjusted p-value, and the other ChAMP output columns. The genes associated with the CpG island with increased or reduced methylation are listed and directly analyzed in EnrichR for transcriptomics results. A summary of these features and the pipeline are shown in **Figure 5**.

**Figure 5.**
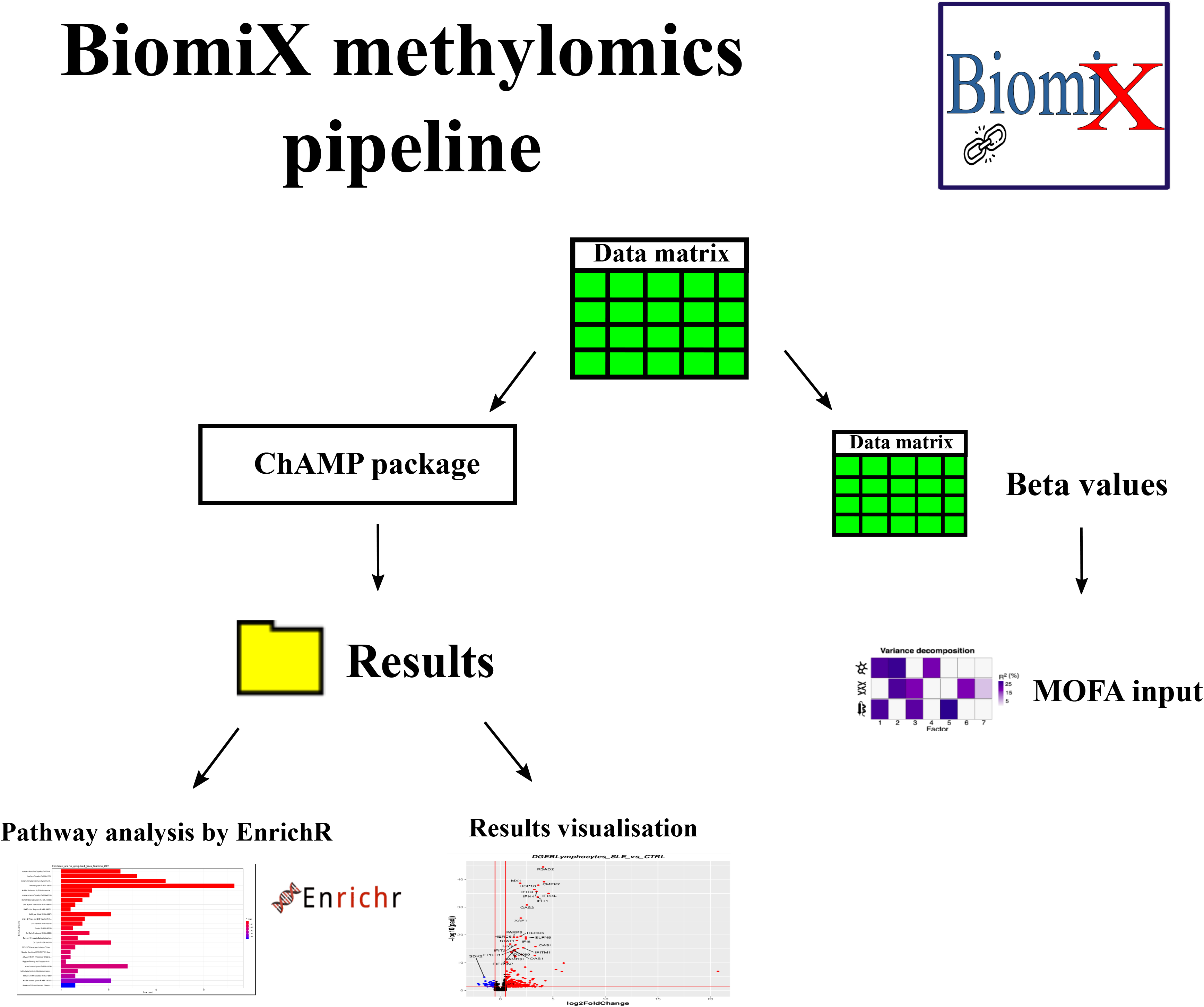
BiomiX Methylomics Pipeline. Illustration of BiomiX methylomics pipeline, showing the beta values input in MOFA integration on the right and statistical analysis on the left. Statistical analysis relies on the ChAMP package analysis, which provides the CpG island with a higher variation between the two groups. Volcano plots and summary files help users explore the results. The biological pathway analysis converts the CpG island to the corresponding gene, if available, and uses it as input. The biological pathway results are available as reports.

### MOFA analysis

MOFA [10] is used according to webpage developer guidelines (https://biofam.github.io/MOFA2/) with normalized input data and reduced feature size. The transcriptomics data are normalized by the variance-stabilizing transformation function in R to make the data approximately homoscedastic. The top genes with higher variance in normalized data are selected for MOFA integration. The metabolomics data are transformed in the pipeline using a log function to improve their Gaussian distribution. All metabolomic peaks are used in the analysis. The methylomics data do not undergo any transformation, but, like transcriptomics, only the top CpG islands with higher variance are selected for integration. MOFA can calculate any desired total number of factors to explain the shared variance between omics datasets. Other parameters customizable in the interface include convergence mode (speed of the convergence), freqELBO (frequence for Evidence Lower Bound Training curve), and Maxiter (number of iteration of MOFA model). The implementation of MOFA in BiomiX includes an automated optimization of the total number of factors. The automatic mode run the MOFA algorithm with an increasing number of factors, stopping the iteration when at least three models shows the last MOFA factor explaining less than 1% of the variability of the data. Only the top three models for separating the two conditions are maintained. How it could discriminate statistically between the two groups is determined for each calculated factor in each model. A non-parametric Mann–Whitney test establishes the factor value distribution between the two groups of samples; the p-values are then corrected using the FDR method. The selected models have the highest number of discriminating MOFA factors. Of the MOFA models with the same number of discriminant factors, only those with the lowest adjusted p-values are selected.

The MOFA analysis provides a matrix containing the variance explained by each factor in a defined MOFA model of n° factors. Two PDFs recapitulate the samples loaded, the variance explained by the factors, and the genes, metabolomic peak signals and/or CpG island contributions of the selected MOFA factor to be explored by scatter plot. Furthermore, a file containing the condition separation performance of each factor and the top 5% of features with an absolute weight of >0.50 (by default but modifiable by the user) are saved as output.

### Extraction of MOFA factors: interpretation

The automatic and non-automatic MOFA analysis includes three methods to ease the user’s interpretation of the discriminating MOFA factors. A summary of MOFA interpretation pipeline in BiomiX is shown in **Figure 6**.

**Figure 6.**
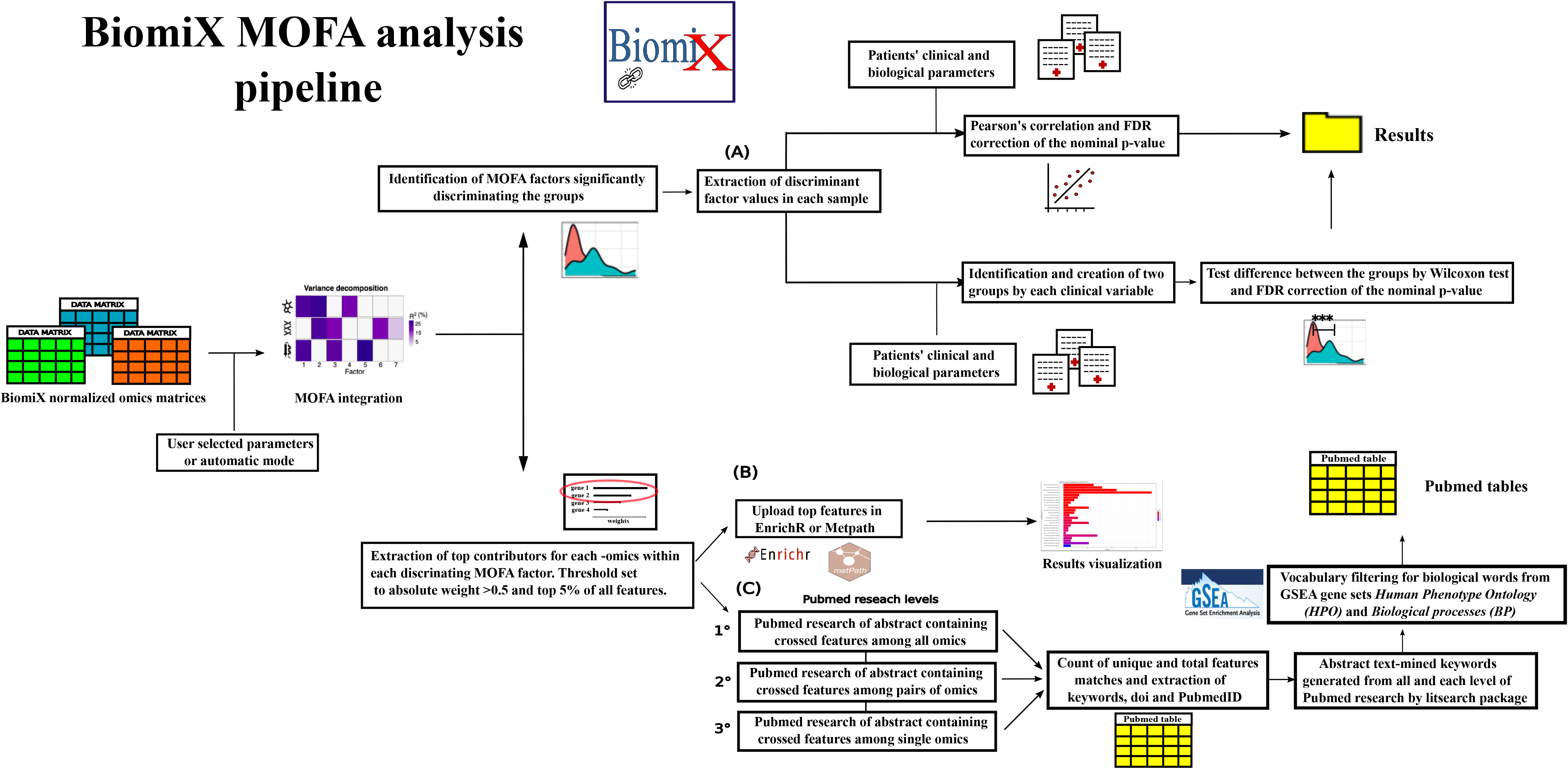
BiomiX MOFA Pipeline. Illustration of the BiomiX MOFA pipeline, showing the input of the different normalized omics matrix to MOFA on the left and its subsequent decomposition in factors. Discriminating MOFA factors and the feature contributing most to them are identified and measured in weights. To simplify the biological interpretation of the factors, BiomiX includes multiple tools to guide users toward a clear understanding of the factors’ nature. The first tool (A) integrates the available clinical and biological data for factor identification. If the clinical data are numeric, they are correlated with each factor in the model by Pearson correlation to spot significant correlations. The nominal p-value is then corrected using the Benjamini–Hochberg method. If the clinical data are labeled, BiomiX forces the identification of two groups, if possible, and parses the samples. The Wilcoxon test infers the importance of that clinical label in the factor, evaluating whether the factor value difference between the two groups is significant. The second tool (B) uses the most contributing features as input for biological pathway analysis, using MetPath and EnrichR, depending on the type of omics. The third tool (C) aims to retrieve PubMed abstracts that have a higher match with the most contributing features of the factor. The BiomiX PubMed search has three levels of research. First, it retrieves abstracts with at least one or more features from all omics. Then, it retrieves abstracts with at least one or more features from each omics pair and, finally, among each single omic. From these three levels, a final table is produced containing the total match and the unique match of features within the abstract, plus the DOI, PubmedID, and keywords. As keywords can be missing in some articles or their number limited, the previously identified abstracts from each level are examined using a text-mining approach in the Litsearch package to extract the keywords. Keyword filtering occurs through a medical–biological vocabulary generated by the terms from GSEA gene sets (Biological process “BP” and Human Phenotype Ontology “HPO”). There is the inclusion of the keywords with higher occurrences at each level, while BiomiX also generates a new document including the match among all the levels.

#### Correlation analysis

Users can upload a matrix containing binary or numerical clinical features to integrate into the MOFA model. The numerical data are correlated through a Pearson correlation with each MOFA factor, while the binary clinical data are analyzed using the Wilcoxon test after dividing the groups into positive and negative categories. The nominal p-values are corrected using the Benjamini–Hochberg method.

#### Pathway analysis

BiomiX retrieves the top contributing genes, metabolites, and CpG islands for discriminating factors in each MOFA model. Depending on the type of omics data, an R package is selected to highlight whether a biological or metabolic pathway is enriched in the enriched genes, metabolites, or CpG islands. The genes are analyzed by EnrichR using the Reactome and biological process, and Encyclopedia of DNA elements (ENCODE) [35] and ChIP-X Enrichment Analysis (ChEA) [36] consensus transcription factors from ChIP-X libraries, while the metabolites are analyzed through MetPath using the KEGG and HMDB databases. CpG islands are associated with their genes, if they exist, and are examined using EnrichR.

#### Pubmed bibliography research

For each discriminating factor in each MOFA model, the top contributing genes, metabolites, and CpG island genes are used as input for PubMed research. The aim is to retrieve the abstracts of articles associated with each discriminating factor to have clues behind their identity for each of them. The search algorithm has three levels of research that prioritize the results of merging more multiomics contributors. Initially, the algorithm selects the top contributors from each omics provided as input, and conducts research in which only abstracts showing at least one out of ten contributors for each omics in the text are selected. The second level does the same, but it selects article abstracts containing at least one out of ten contributors in omics pairs (e.g., transcriptomics–metabolomics, methylomics–transcriptomics, and metabolomics–methylomics). Finally, the last-level research selects article abstracts showing at least one out of ten contributors within a single omics in the text.

For these three levels, the output document includes a .TSV table containing the PubMed articles, the total number of matches among the total number of contributors and the number of times contributors. Keywords, DOIs, and match information for each contributor are available. The author-provided keywords are not optimal due to their absence in some journals. Therefore, BiomiX includes further text-mining analysis. The article’s abstract, spotted at each level, is extracted and parsed through litsearchr version 1.0.0 in a two-to-four-word combination [37]. The vocabulary generated by each abstract is analyzed to identify the more frequent combinations of words. These words are filtered by another vocabulary comprising gene set names in Gene Ontology biological processes (7,751 gene sets) and human phenotype ontology (5,405 gene sets). The 15 most frequently used words are included in the output .TSV file. Finally, the analysis of the abstracts retrieved at all three levels is repeated.

### Case studies

Two multi-omics datasets were used to test BiomiX. First, the FastQ of tuberculosis dataset were downloaded from ENA (https://www.ebi.ac.uk/ena/browser/home) with the project code PRJNA971365. The quality was checked by FastQC and trimmed by Trimmomatic v0.39. The FastQ files were aligned to the Ensembl Homo sapiens reference genome (GRCh38) and annotated to GENCODE GRCh38.104 using STAR v2.7.11 running two-pass mapping strategy with default parameters. Gene quantification was performed using Ht-seq count v0.13.538 default parameters. At the end of the process, the 13 samples per each condition, *i.e.* healthy controls (HC), patient with tuberculosis (PTB) and patient with tuberculosis and diabetes (PTB_DM) were available. For the sake of clarity, only the comparison between HC and PTB is reported. The entire dataset is available as example dataset in BiomiX. The parameters were set to reproduce those similar to the original work, including |log2FC| > 1 and p.adj < 0.05 for the transcriptomics analysis, and |log2FC| > 0.5 and p.adj < 0.1 for the metabolomics analysis. Second, the FastQ of Chronic Lymphocytic Leukemia (CLL) dataset were downloaded from http://pace.embl.de/ and analyzed according to |log2FC| > 1 and p.adj < 0.05 threshold for the transcriptomics analysis, and |log2FC| > 0.5 and p.adj < 0.1 threshold for the metabolomics analysis.

## Results

### Implementations and comparison with other tools

BiomiX aims to provide a user-friendly tool connected to other well-established platforms, such as Metaboanalyst, EnrichR and GSEA. BiomiX uses matrix-compliant data formats available in the main public repositories such as Sequence Read Archive (SRA) and Metabolight to support the reproducibility and reusability of the workflow, in line with the FAIR (Findable, Accessible, Interoperable, and Reusable) principle. Specifically, for metabolomics, the metabolite annotation/assignment files (MAFs) can provide the remaining information about annotation. Fragmentation .mzml files can be used for the MS2 annotation. Finally, the BiomiX-assisted format convertor can fix compatibility issues by converting the matrix in the proper format with a simple user interface. BiomiX is the first tool to analyse single omics and integrate them with MOFA integration. Moreover, BiomiX has also implemented the MOFA algorithm to select the models containing factors that discriminate between the two conditions of interest. The advantage of this unsupervised approach is the identification of differences between groups based on the unbiased nature of the analysis, which highlights only group-independent common omics changes. The identification of a group-discriminant factor in this context must account for a considerable explained variance, proving its reliability and palpability. A BiomiX breakthrough is the multi-approach adopted in the identification of the MOFA factor, in particular, the innovative bibliography search on Pubmed carried out on the contributors of the significant factors, a novel approach that can lead to the interpretation of factor together with the analysis of biological pathways on the contributors and the correlation analysis of the significant MOFA factors with the available clinical data. BiomiX approach differs from X-omics ACTION [38], the novel Nextflow [39] pipeline implementations for multiomics analysis, which relies solely on simple correlation analysis with clinical data for MOFA interpretation. Moreover, X-omics ACTION integrates only metabolomics and methylomics data, without analyzing them. This is an important limitation given the wide diffusion of transcriptomics data. The MixOmics project was and still is a milestone in multi-omics integration but it currently only provides supervised integration approaches (DIABLO) for multi-omics data from the same patient, without analyzing single omics data separately. Other tools that do not integrate omics data but focus on single omics analysis are iDEP [40] and Metaboanalyst [9], for transcriptomics and metabolomics, respectively. These tools provide a wide range of downstream analyses such as heatmaps, PCA, pathways, and network analysis to facilitate data exploration. Similar functionalities are provided in BiomiX exploiting the same packages. To enable the use of functionalities not included (PCA or network analysis), BiomiX generates output files that can be processed by other user-friendly programs. BiomiX is not a stand-alone tool but can connect to existing tools in the single omics analysis and goes beyond providing access to the multi-omics integration of these data. Compared with bioinformatics tools specifically designed for the integration of a single omics or multi-omics (**Table 1**), BiomiX’s features bridge a gap in the lack of multi-omics analysis and integration tools, highlighting its importance.

**Table 1.**
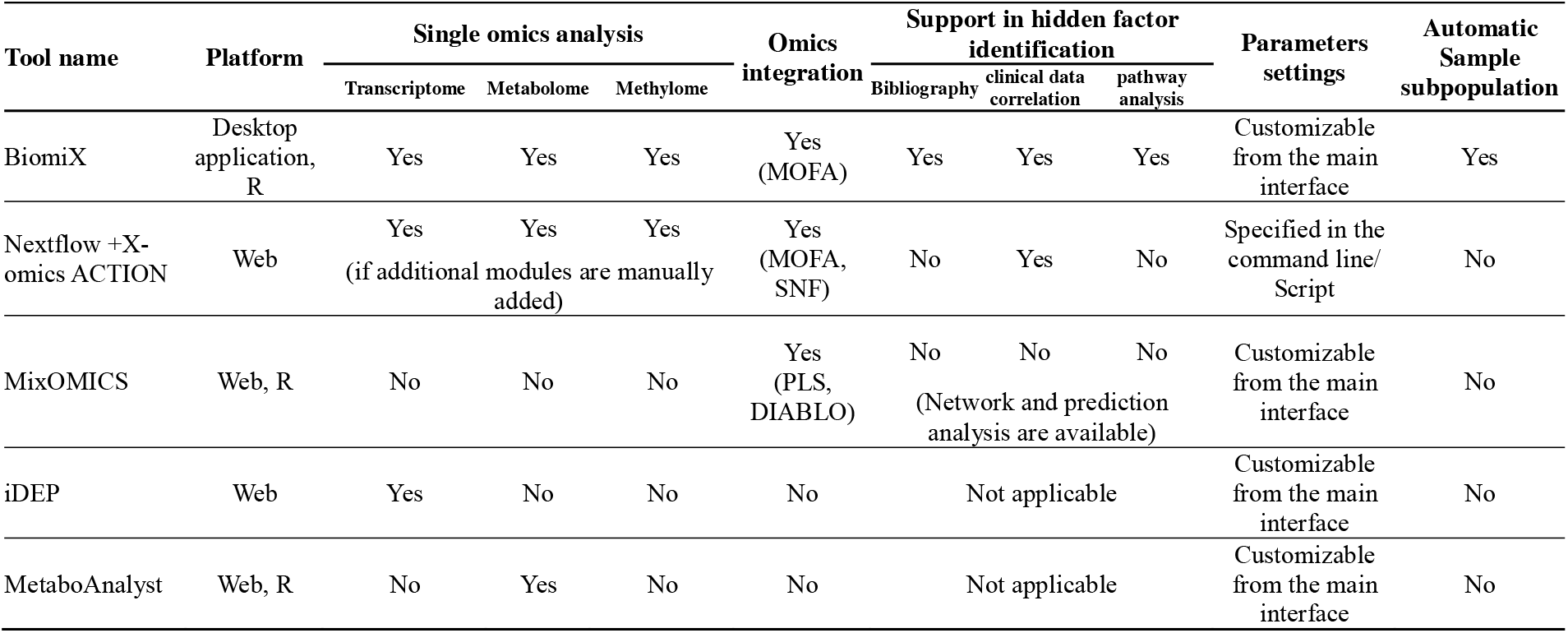
Comparison of BiomiX with single omics and multi omics integration tools.

### Case studies

BiomiX has been applied to two example of databases composed of multiomics data. The first case study included a transcriptomics and metabolomics analysis on PTB [41]. Compared to CTRLs, the transcriptomics analysis of BiomiX showed the same biological pathways as in the manuscript, including Arachidonic Acid Metabolic Process (GO:0019369), Classical Antibody-Mediated Complement Activation (R-HSA-173623), but also novel ones as Hydrolysis Of LPC (R-HSA-1483115) and B Cell Receptor Signaling Pathway (GO:0050853). The pathways were consistent in both the Gene Ontology and Reactome databases. Furthermore, to assess differences in patients’ immune responses, BiomiX enabled patients to be subgrouped according to a gene panel containing 26 IFN-induced genes. All samples having at least one IFN gene with a Z-score > 1 were considered positive, providing a clear separation (**Figure S1**). At first all the PTB transcriptome was compared with the HC (**Figure S2A and B**). The IFN negative subgroup had genes differentially expressed enriched in Nitric Oxide Biosynthetic Process (GO:0006809) in response to infection [42], but also Arachidonic Acid Metabolic Process (R-HSA-2142753), suggesting an IFN-independent activation. IFN positive were enriched in IFN signaling R-HSA-877300, B Cell Receptor Signaling Pathway (GO:0050853) with a reduction of IL-10 production (R-HSA-6783783) as expected. The metabolomics analysis on plasma revealed reductions in Acylcarnitine, PC, LysoPE and TG, with significant enrichment for the Sphingolipid signaling pathway, Retrograde endocannabinoid signaling, caffeine, purine, linoleic and Glycerophospholipid metabolism as in the manuscript, but also novel ones as Phenylalanine metabolism and Fc gamma R-mediated phagocytosis (**Figure S3A and B**). Data integration was then carried out by BiomiX implementation of MOFA integration, which consists of two steps. First, the iterative calculus of MOFA models with a progressive number of total factors in each iteration stopped when at least three models had the last factor variance explaining less than 1%. Next, the three best-performing models in separating the two conditions by the Wilcoxon test were selected, based on the number of discriminant factors and the adjusted p.values (p.adj). Here, the MOFA factors 3, 4 and 5 were selected, with the first offering the best separation between PTB and HC and therefore being used for the following analysis. The three-factor MOFA models explained the 5.05% and 45.77% metabolomics and transcriptomics total variances, respectively. Furthermore, only factor 1 significantly discriminated the two conditions (p.adj = 0.0011, sd = 0.08) (**Figure 7A and B**), catching 2.23% and 33.22% of metabolomics and transcriptomics total variances, respectively, but its identity needs identification. BiomiX provided this interpretation of factor 1 through its innovative bibliography search, which by evaluating the transcriptomic contributors to factor 1, spotted articles related to inflammation-driven genes, bacterial inflammation and even more specifically, articles related to Mycobacterium tuberculosis infections. The metabolomic contributors also suggested two articles related to bacteria but less specific for inflammation and less informative. Pathway analyses of the positive contributors of factor 1 confirmed the inflammation and interferon as the main biological process (R-HSA-913531, R-HSA-1280215, R-HSA-1169410) while, only Trilostane, an inhibitor of corticoid production, was identified in metabolomics positive contributors. Consistently, the negative transcriptomics contributors Interleukin-10 Signaling and immunoregulatory pathways were enriched (R-HSA-6783783, R-HSA-198933), while 11-deoxycortisol, PC, Sphingolipids and cholesterol were the main negative metabolomics contributors, enriched in Sphingolipid Metabolism and Alpha Linolenic Acid and Linoleic Acid Metabolism (p-value < 0.05). The inflammation-induced lipid alteration known in the literature [43], and the negative contribution of anti-inflammatory factors (IL-10 and corticoids) support the inflammation-interferon identity of factor 1. The results obtained from BiomiX are available in the **Supplementary table 1**.

**Figure 7.**
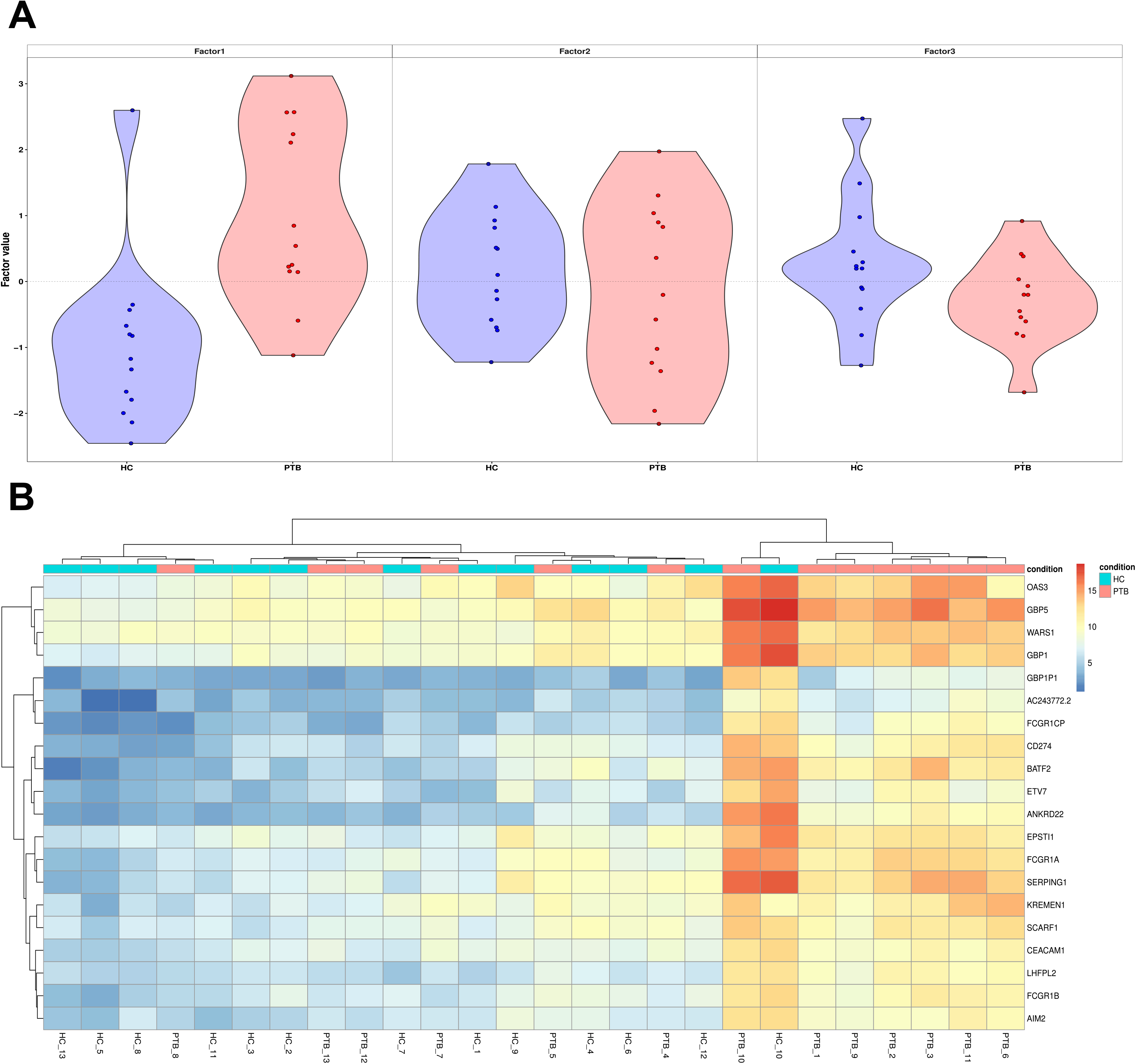
Whole blood transcriptomics and plasma metabolomics analysis from tuberculosis patients using BiomiX. **A)** Violin plot representing the distribution of patients affected by Tuberculosis (PTB; red) and control samples (HC; blue) based on each MOFA factor’s value. **B)** Heatmaps using the top whole-blood 20 genes contributing to Factor 1. PTB patients (red squares) and HC (blue squares) are shown on the top. Heatmap distance: “Euclidean”, clustering method “complete”. The gene expression was normalised using the variance stabilising transformation (VST) method.

The second case studied the transcriptomics and methylomics differences in CLL patients with mutated and unmutated IGHV. This dataset has been used for testing the MOFA algorithm. Data presented here will thus mainly focus on the identification of the identity behind the factors, briefly citing the transcriptomics and methylomics single omics results. Transcriptomics (**Figure S4A and B**) and methylomics heatmap and volcano plot (**Figure S5A and B**) successfully highlighted the genes related to CLL differences mentioned in the article (KANK2, DGKH, MYLK, PPP1R9A, SEPTIN10, SOWAHC, PLD1, and LPL). Moreover, the pathway analysis identified upregulation in proliferation (GO:0090267) and VDJ recombination (GO:0033152, GO:0033151) pathways in methylomics. The MOFA implementation identified the model with 8, 9 and 10 factors as the best performing for identifying differences between the two conditions. The ten-factor model was the best, identifying five factors (factors 1, 3, 4, 5 and 6) discriminating mutated and unmutated IGHV CLLs (**Figure 8A**). Factor 1 was the most discriminating among them, and its analysis (**Figure 8B**) identified articles related to DNA damage, ageing and cancer. Correlation analysis identified a significant correlation with 11q22.3 deletion, known to induce proliferation in CLL by ZAP-70. These results confirmed the proliferation and oxidative processes already identified by the MOFA article. Factor 3 and factor 4 were correlated with chromosome 12 trisomy, while the bibliography search identified articles related to B leukemia, DNA damage, and autoimmunity specifically for factor 3. Factor 4 was enriched in focal adhesion and membrane-ECM Interactions and signaling (R-HSA-3000171, R-HSA-8874081), consistent with the roles of trisomy 12 in activating adhesion signaling [44]. Factor 5 was correlated with gender differences in CLL (p.adj = 1.78e-26) in accordance with articles related to sex differences in human-primates [45], mice sex development [46] and autoimmune encephalomyelitis [47]. Furthermore, it is the only factor not correlated to any treatment response. Interestingly, the pathway analysis highlighted pathways linked to galactosyltransferase activity (R-HSA-4420332, R-HSA-3560801, R-HSA-3560783), suggesting a sex difference in CLL that is not targeted by any drug. Finally, factor 6 was associated with mutations (TP53, SF3B1) and a deletion (del17p13) associated with a poor outcome and unfavorable prognostic factors [48,49]. The pathway analysis supported this hypothesis, as contributors were enriched for cytokine signaling such as interleukin 4, 13, 27 (GO:0070106, R-HSA-6785807) and type 1 interferon (GO:0032481), of which interleukin 4 and IFN-alpha are already known to be linked to a more severe condition [50]. The results obtained from BiomiX are available in the **supplementary table 2**.

**Figure 8.**
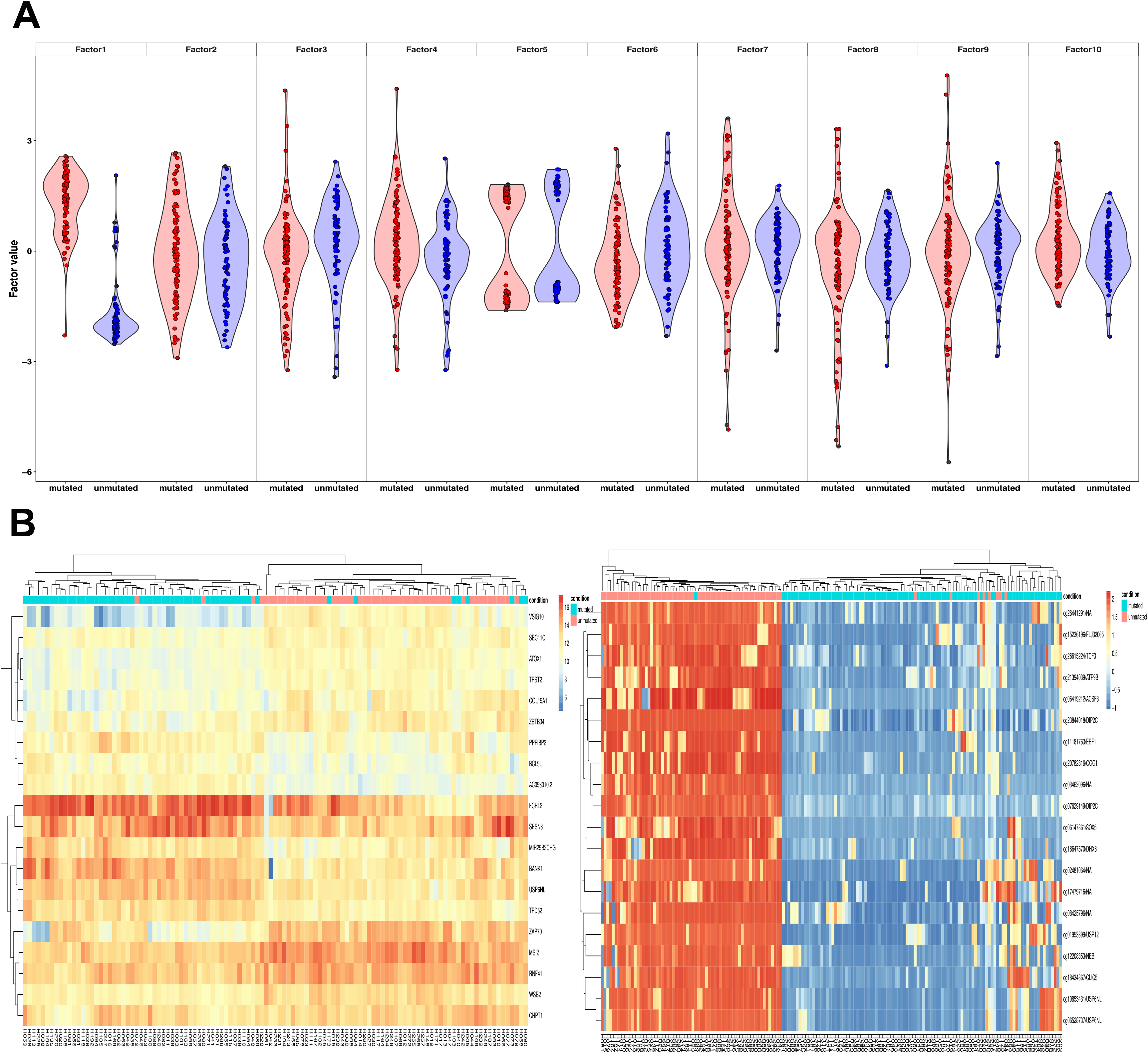
Whole blood transcriptomics and plasma metabolomics analysis from chronic lymphocytic leukemia patients using BiomiX. **A)** Violin plot representing the distribution of patients affected by chronic lymphocytic leukemia (CLL) with mutated (red) and unmutated (blue) IGHV based on each MOFA factor’s value. **B)** Heatmaps using the top whole-blood 20 genes contributing to Factor 1, showing whole blood transcriptome (left) and whole-blood methylome (right). IGHV unmutated CLL (blue squares) and IGHV mutated CLL (red squares) are shown on the top of the heatmaps. Heatmap distance: “Euclidean”, clustering method “complete”. The gene expression was normalised using the variance stabilising transformation (VST) method.

### Discussion

BiomiX aims to help biologists, physicians, and scientists with no background in bioinformatics. Some tools, such as MixOmics, allow for data integration, while others perform single-omics analysis, but none of them allow for both simultaneously [51]. Currently, to our knowledge, the only way to compare two groups with an analysis similar to that of BiomiX, is the Nextflow association of a single omics pipeline (metaboigniter [52] or other transcriptomics and methylomics modules) with integration tools such as X-omics ACTION. However, this pipeline does not contain implementations such as the MOFA number of factors optimization. Moreover, while X-omics ACTION only performs correlation analysis to annotate MOFA factors, BiomiX relies on three methods based on different approaches that converge to reveal the factor’s identity, as shown in the two case studies. BiomiX is thus more precise in providing factor annotation and is the first tool to exploit an innovative Pubmed bibliography to annotate hidden factors. It is also worth mentioning that although Nextflow has developed a user interface, most changes to the pipeline require coding skills. BiomiX solution does not require coding skills and is not definitive; rather, it is a first attempt to incorporate the gold standard single-omics pipeline in bioinformatics with automated integration using MOFA. BiomiX guarantees high interpretability of the common source of variation among omics, providing users with single results from omic and multiomics integration. It allows for a complete overview of the changes occurring in biological systems, either in the context of disease, treatment, or physiological conditions, enhancing the interpretability of the biological pathways and processes involved. In addition to improving MOFA factor interpretability and optimizing the total number of factors, BiomiX creates a user-friendly interface. BiomiX also provides output formats ready to copy and paste into specialized user-friendly widespread websites or programs, such as GSEA, EnrichR, and Metaboanalyst. Nevertheless, BiomiX has limitations. Although it can analyze multiple groups simultaneously, it has limited types of omics analyzers. Consequently, much work remains to be done to implement functionalities based on community needs and to include more integration methods and omics data, such as proteomics and genomics data from different technologies. We aim to grow the community of users and developers to improve the accessibility of new bioinformatics algorithms and methods within the scientific community. BiomiX represents this attempt to improve accessibility to bioinformatics tools by offering everyone access to bioinformatics methods and algorithms, enabling specialists in a wide range of fields to benefit from the Big Data revolution.

## Author Contributions

CI was in charge of writing and planning the bioinformatic approaches. CJ and AB contributed equally to supervising the work and the scientific relevance of the article, while the other authors evaluated, revised, and approved the article.

## Fundings

This work was supported by the Innovative Medicines Initiative Joint Undertaking under Grant Agreement Number 115565, resources of which are composed of financial contributions from the European Union’s Seventh Framework Program (FP7/2007–2013) and EFPIA companies in kind.

## Acknowledgments

We also acknowledge the 3TR and PRECISESADS consortium, which guaranteed the data used in this article and allowed us to analyze them.

## Code Availability

The download and tutorial are available on the following BiomiX github pages, respectively: https://github.com/IxI-97/BiomiX2.2 and https://ixi-97.github.io.

**Figure S1. Identification of IFN-positive and negative patients affected by tubercolosis (PTB) with BiomiX.** The classification into positive and negative samples was obtained by hierarchical clustering based on the Z-score measured for each gene within each sample. A single square corresponds to the Z-score of a specific IFN gene for an individual sample. A single IFN gene having Z-score > 1 was enough to include a patient in the positive group. Euclidean distance and Ward’s D2 method have been used for hierarchical clustering.

**Figure S2. Identification of patients affected by tubercolosis (PTB) based on transcriptomics dataset with BiomiX. A)** Volcano plot of the differential gene expression (DGE) analysis of PTB samples vs healthy control samples (HC) in the whole blood dataset. Downregulated and upregulated genes are depicted in blue and red, respectively. The top 15 genes in each category are indicated. The significance thresholds shown as red lines are |Log2FC| > 1 and false discovery rate (FDR) < 0.05. **B)** Heatmap of the top 20 upregulated genes and top 20 downregulated genes. Genes are depicted in the rows while each column represents a sample (HC in blue and PTB in red on the top). The genes were normalized by vst method. Euclidean distance and Ward’s D2 method have been used for hierarchical clustering.

**Figure S3. Identification of patients affected by tubercolosis (PTB) based on plasma metabolomics dataset with BiomiX. A)** Volcano plot of the metabolite comparison of PTB samples vs healthy controls (HC) in the plasma metabolomics dataset. Downregulated and upregulated metabolites are depicted in blue and red, respectively. The top 15 metabolites in each category are indicated. The significance thresholds shown as red lines are |Log2FC| > 1 and false discovery rate (FDR) < 0.05. **B)** Heatmap of the top 20 upregulated metabolites and top 20 downregulated metabolites. Metabolites are depicted in the rows while each column represents a sample (HC in blue and PTB in red on the top). The metabolite peaks were log transformed. Euclidean distance and Ward’s D2 method have been used for hierarchical clustering.

**Figure S4: Identification of chronic lymphocytic leukemia (CLL) patients based on transcriptomics dataset with BiomiX. A)** Volcano plot of the differential gene expression (DGE) analysis of mutated IGHV CLL samples vs unmutated IGHV CLL samples in the whole blood dataset. Downregulated and upregulated genes are depicted in blue and red, respectively. The top 15 genes in each category are indicated. The significance thresholds shown as red lines are |Log2FC| > 1 and false discovery rate (FDR) < 0.05. **B)** Heatmap of the top 20 upregulated genes and top 20 downregulated genes. Genes are depicted in the rows while each column represents a sample (unmutated IGHV CLL in blue and mutated IGHV CLL in red on the top). The genes were normalized by vst method. Euclidean distance and Ward’s D2 method have been used for hierarchical clustering.

**Figure S5. Identification of chronic lymphocytic leukemia (CLL) patients based on methylomics dataset with BiomiX. A)** Volcano plot of the differential methylation analysis of mutated IGHV CLL samples vs unmutated IGHV CLL samples in the whole blood dataset. Downmethylated and upmethylated genes are depicted in blue and red, respectively. The top 15 genes in each category are indicated. The significance thresholds shown as red lines are |Log2FC| > 1 and false discovery rate (FDR) < 0.05. **B)** Heatmap of the top 20 upmethylated genes and top 20 downmethylated genes. Genes are depicted in the rows while each column represents a sample (unmutated IGHV CLL in blue and mutated IGHV CLL in red on the top). Euclidean distance and Ward’s D2 method have been used for hierarchical clustering.

## Notes

https://github.com/IxI-97/BiomiX2.2

https://ixi-97.github.io

### Competing Interest Statement

The authors have declared no competing interest.

