## Supplementary figures and images for "BiomiX, a User-Friendly Bioinformatic Tool for Automatized Multiomics Data Analysis and Integration"

### Figure_S1.jpg

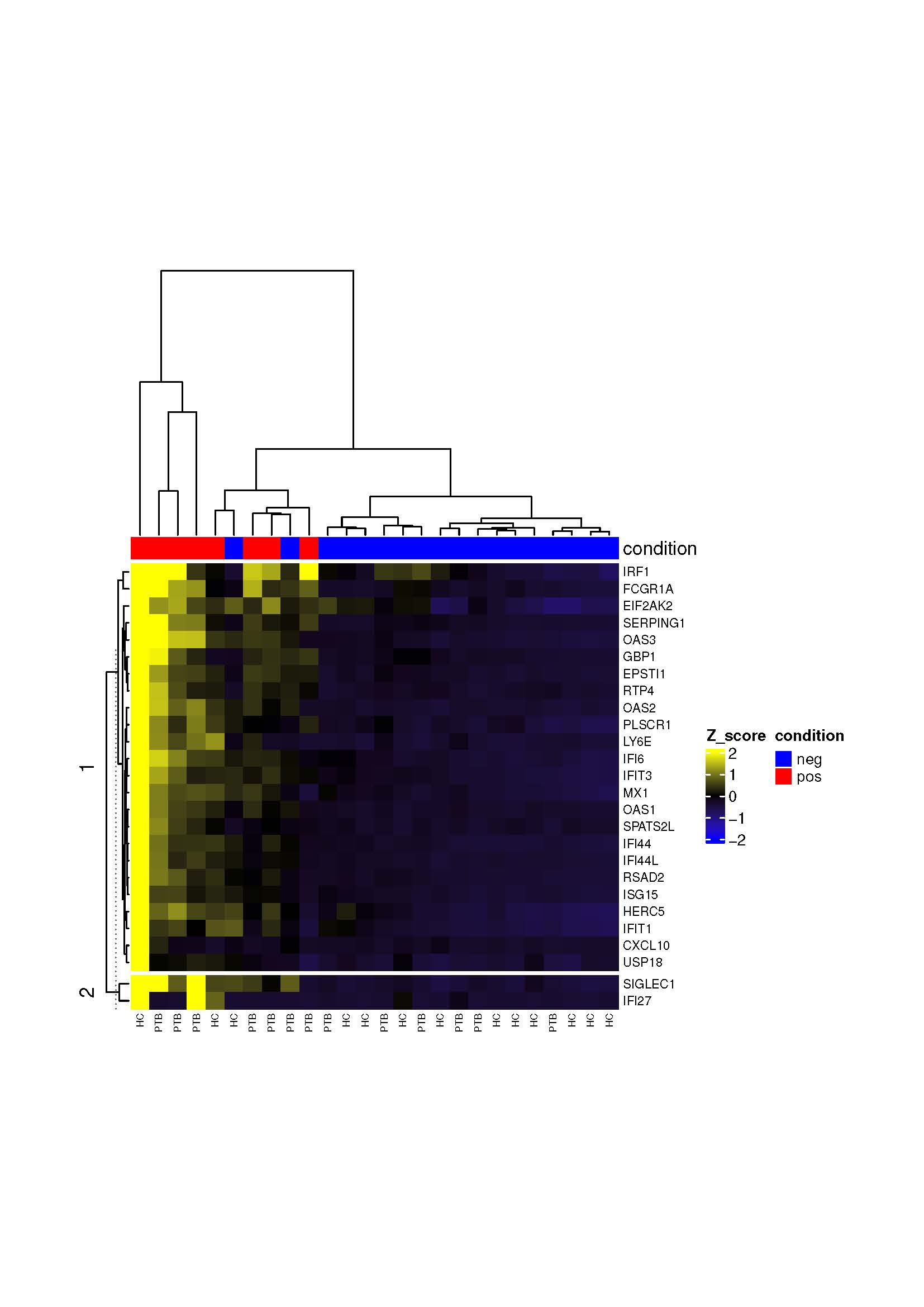
